# Sex-specific roles for yawning in the Emperor cichlid *Boulengerochromis microlepis*

**DOI:** 10.64898/2026.05.28.728490

**Authors:** Ayasha Abdalla-Wyse, Jaden Y. Da Silva, Ritambhara Kalia, Vasundhara Kalia, Samruddhi Landge, Purvi R. S. Munasinghe, Casey C. Schwartz, Maxwell E. R. Shafer

## Abstract

Yawning is a poorly understood, yet spontaneous and ubiquitous behaviour. Three main hypotheses explain the function of yawns, most notably stimulating arousal. Yawns can also play a role in social cohesion through their “contagious” nature, eliciting an unconscious yawn in an observer. Yawn contagion has been observed in mammals and birds, and has been suggested in fish. Yawning also contributes to sexual dimorphisms, whereby males and females display differences in yawn frequency or function. Here, we examine these hypotheses in juveniles of the African cichlid, *Boulengerochromis microlepis*. By re-analyzing hundreds of hours of video recordings and nearly 3000 observed yawns we find support for all three yawn hypotheses in fish and two temporally and sexually distinct effects for yawns. First, our results suggest yawns at night and dawn induce arousal by delaying transitions into sleep. In contrast, daytime yawns induced rapid yawn contagion in neighbouring fish. Our analyses also demonstrate clear differences between sexes, with increased yawning in males at night and dawn associated with less sleep, whereas females displayed more robust social yawning. Together, our results expand the evolutionary context for yawning, and raise further questions about its functions in sleep and social cohesion across animals, and across sexes.

## INTRODUCTION

Yawning is a spontaneous and uncontrollable behaviour that has been observed across the animal kingdom and likely arose with the evolution of jawed fish^1^. Despite being a widespread and well studied phenomenon, the function of yawning is still poorly understood. Many theories exist for yawning, including roles in the modulation of sleep/wake transitions and arousal, induction of social cohesion through contagious yawning, and contributions to sexual dimorphisms within species (yawn frequency or functions). Though yawning has been observed in all jawed animals including fish, these theories have primarily been thoroughly tested in mammalian species.

Yawning is associated with many physiological, social, and neurological factors across jawed animals. One of the strongest connections to yawns is their relation to the sleep-wake cycle. Yawns are linked to the circadian rhythm, where humans show a bimodal distribution, with yawns occurring in the morning after waking up and at night before going to sleep^1,2^. The state change hypothesis is a long-standing idea that yawning occurs before or after a behavioural state change, such as wake to sleep or sleep to wake^3^. This has been observed across the animal kingdom, in both endotherms, including primates^4,5^, lions (*Panthera leo*)^6^, and dolphins (*Tursiops truncatus*)^7^, as well as ectoderms, like salamanders (*Phaeognathus hubrichti*)^8^ and fish (char, *Salvelinus leucomaenis*)^9^. While it is known that yawns are often accompanied with physiological changes associated with increased arousal^10^, it is unclear whether yawns are directly impacting transitions between sleep and wake states in all species.

Sex differences have also been observed for yawns in some species. The sexual dimorphism hypothesis stems from the observation that males of highly dimorphic primate species tend to yawn more than females. This has been observed in chimpanzees (*Pan troglodytes*)^11^, the grey-cheeked mangabey (*Cercocebus albigena*)^12^, and some macaques (*Macaca fascicularis*^12^, *M. nigra*^13^ and *M. fuscata*^14^). Interestingly, this is only seen in primates for which sexual dimorphism is characterized by canine size; ring-tailed lemurs (*Lemur catta*), Verreaux’s sifaka (*Propithecus verreauxi*)^4^ and humans (*Homo sapiens*)^15,16^ do not show sex differences in canine size or yawn frequency. Evidence for sexual dimorphism in yawning is also observed in some non-primate mammals that show sexual dimorphism in other sex characteristics, such as sea lions (*Otaria flavescens*)^17^ and rats^18,19^, where males yawn more than females. This type of yawn may be an aggressive or threat display, for dominant males to display their canines or mouth size, which is an indicator of social status^4^. However, it is unclear if this phenomenon is limited to mammals.

One of the most fascinating characteristics of yawning is its contagious nature. Many studies within different animal groups have shown that observing a yawn can produce a yawn in an observer of the same species or even in intra-species pairs. Yawn contagion is so potent that even artificial yawns (video recordings)^20^, yawn sounds^21^, or even just the thought of yawns can elicit a yawn response in many different species, including humans^3^. In addition to humans and other primates, yawn contagion has been consistently shown in non-primate mammals, including wolves (*Canus lupus lupus*)^22^, dogs (*Canis lupus familiaris*)^23^, pigs (*Sus scrofa*)^24^, African elephants (*Loxodonta africana*)^25^, and sheep (*Ovis aries*). These observations extend outside of mammals, including in social birds like budgerigars (*Melopsittacus undulates*)^26^, and even evidence of contagion in zebrafish (*Danio rerio*)^27^. Generally, yawns are considered contagious for up to 5 minutes after the original stimuli, with most contagious responses occurring within 1 minute^3,28^. Current hypotheses suggest that the contagious aspect of yawning emerged recently in species with complex social hierarchies, playing roles in communication, and to indicate tiredness, boredom, or social tension^1^. However, it is unclear when this behaviour evolved or if it is unique to social animals or specific clades (e.g. mammals or birds).

The emperor cichlid, *Boulengerochromis microlepis,* is the largest species within the Lake Tanganyikan adaptive radiation of cichlid fish^29,30^. Cichlids display extensive morphological, ecological and behavioural diversity, including an array of different complex sleep^31,32^ and social behaviours^33,34^. Here, by re-analysing video recordings and live tracks of freely behaving and interacting diurnal *B. microlepis* individuals, we test multiple hypotheses for the function of yawning. We examined the temporal distribution of yawns and determined the time between yawns and changes between wake and sleep states to test the state change hypothesis. By comparing the time of yawns between neighbouring individuals, we confirm the existence of yawn contagion in fish. We find evidence to support the sexual dimorphism hypothesis in fish, observing differences in the temporal distribution and function of yawns between males and females. We find support for the state change hypothesis, and that yawns stimulate arousal around behavioural transitions. We then speculate on the roles of yawning and yawn contagion across animals.

## RESULTS

### *B. microlepis* show sex differences in the temporal frequency of yawns

In a previous study, the activity of multiple freely behaving individuals of *B. microlepis* were recorded, and wake (active) and rest states were identified based on swimming speed and a rolling window threshold for activity^31^. In these data, rest is used as a proxy for sleep, and is analogous to similar states in zebrafish and other fish species that meet the behavioural criteria for sleep^35^. In analyzing these recordings, we observed that individual fish yawned frequently, opening their mouths fully and stretching their fins, analogous to yawn behaviours across animals, including humans (**Fig. 1a**). Taking advantage of our high resolution and extensive recordings, in the present study we quantify and analyze these yawns to test three theories for the existence and functions of yawns in animals (**Fig. 1b**). First, we examine if there are differences in the frequency, timing, or functions of yawns between females and males. Second, we examine the roles of yawns in state transitions between active and inactive (sleep) states. Third, we test if yawns are contagious in cichlid fishes.

**Figure 1.**
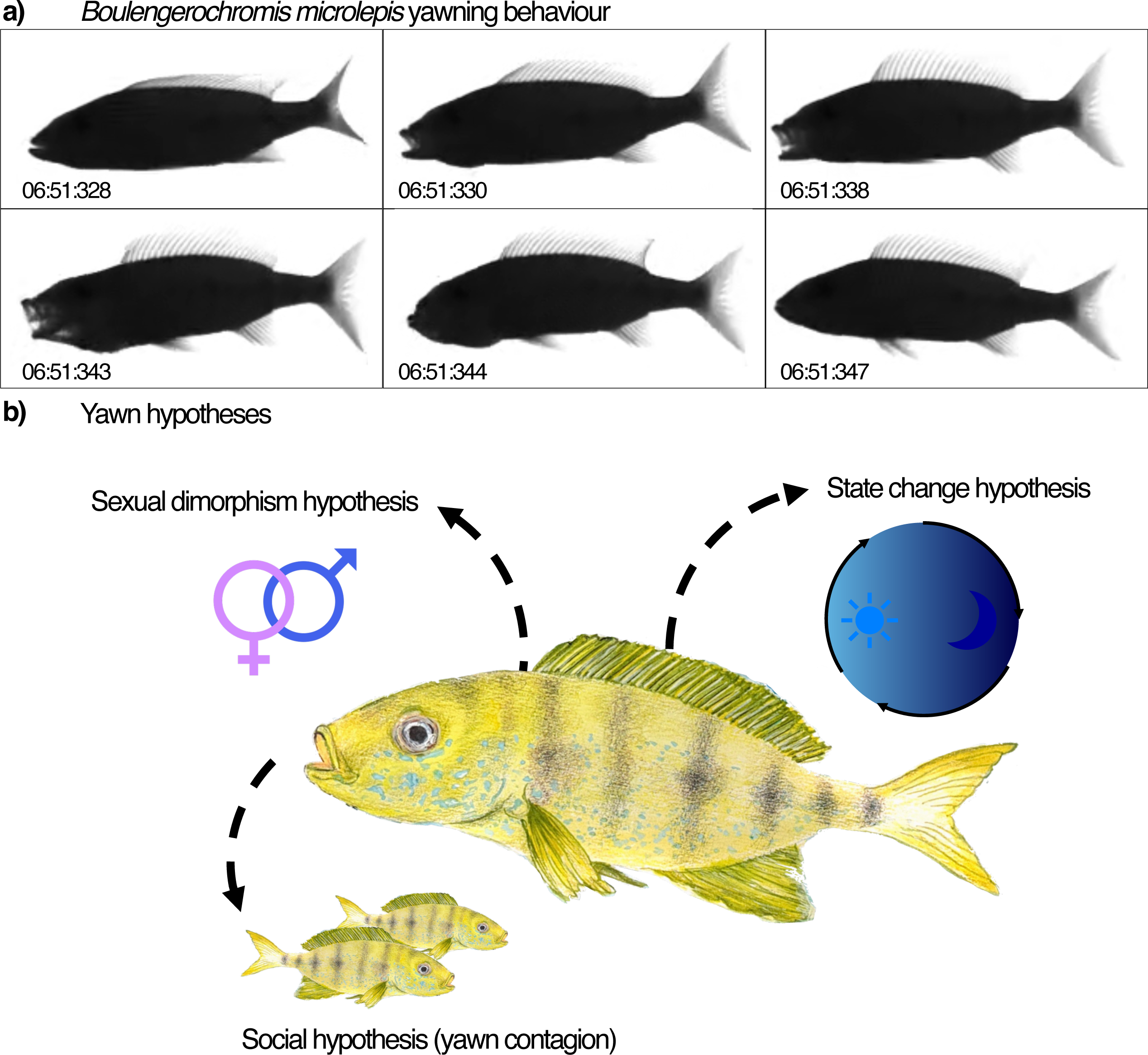
Overview of yawn behaviours and hypotheses tested in *B. microlepis*. **a)** Image stills from video recording of individual ID FISH20200923_c3_r0, from 06:51:328 to 06:51:347 in the morning (1.9 seconds long), depicting the stages of a yawn, in which the fish fully opens and stretches its jaw and fins, then closes their mouth. **b)** Yawn hypotheses that were tested in *B. microlepis*. Illustrations by Vasundhara Kalia.

A total of 2917 yawns were observed over a 48 hour (h) period across seven individuals and > 300 hours of recordings (**Fig. 2a, Supplemental Data 1**), with a mean of 416.71 yawns per individual (± 93 SD). No differences were observed in the total amount (*P* = 0.20) or duration (*P* = 0.67) of yawns between males (*N* = 3) and females (*N* = 4) (**Fig. 2b-c**). However, we did observe a difference in the temporal distribution of yawning between sexes (**Fig. 2d**), whereby males yawn at a significantly higher rate at night and dawn (13.75 - 12.5 yawns per hour) compared to during the day or dusk (6.55 - 4.08 yawns per hour; *P* = 0.0037), and females show a steady rate of yawns per hour in every phase (between 5.38 - 8.36 yawns per hour; *P* = 0.21). These results suggest that these fish display sexual dimorphism in the timing of yawns, but not the total amount or duration of yawns per individual.

**Figure 2.**
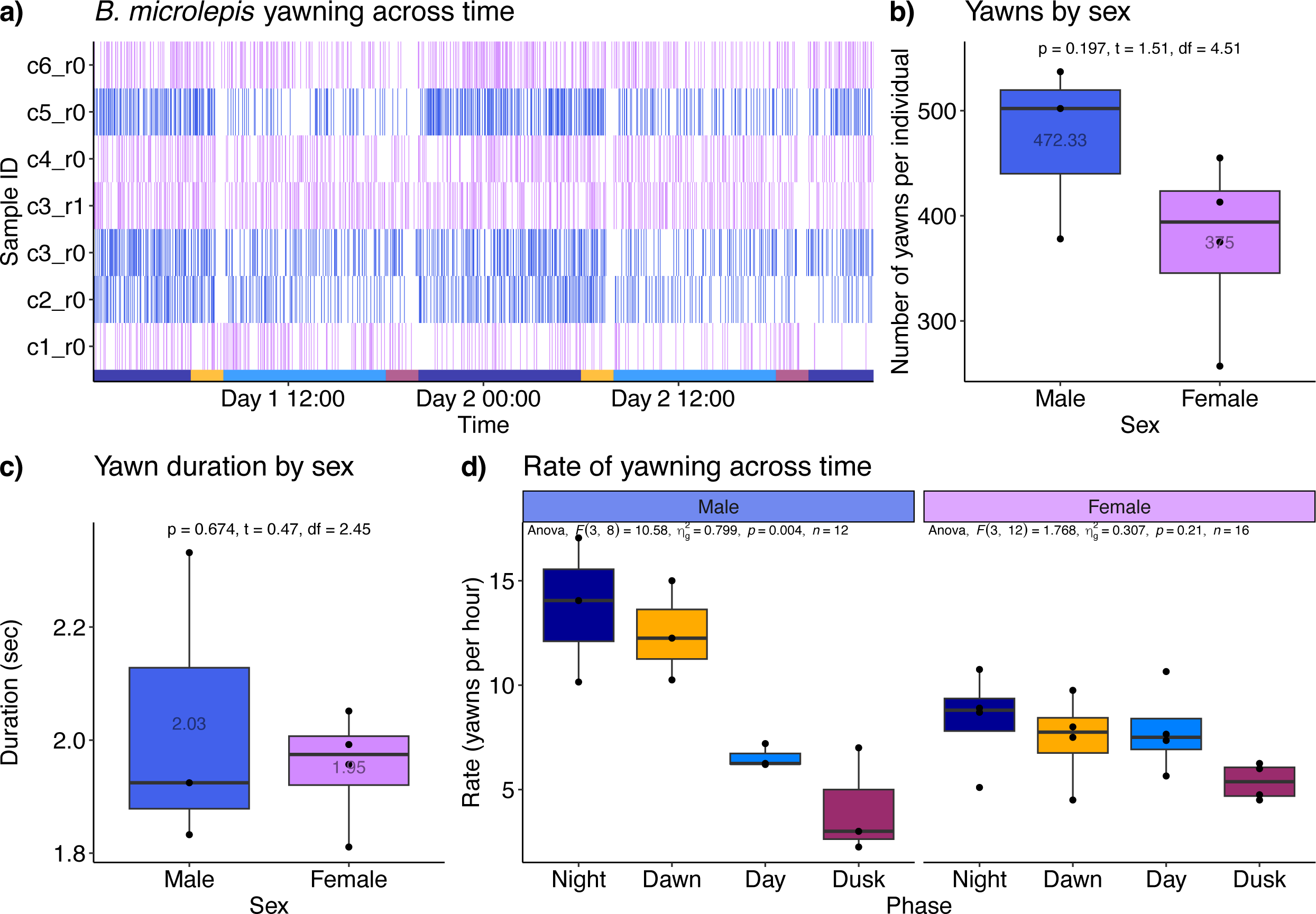
*B. microlepis* yawns are circadian. **a)** Raster plot showing the timing of 2917 yawns for all samples tested across 48 hours. **b-c)** Boxplot showing the number of yawns per individual (**b**) and duration of yawns (**c**) between sexes across individuals of *B. microlepis*. An unpaired t-test was used to compare sexes. **d)** Boxplot of the rate of yawning (yawns per hour) across time in *B. microlepis*. An analysis of variance (ANOVA) was used to compare the rate of yawning within sexes across time between the four phases. Boxes represent the interquartile range (IQR) from the first to third quartile, where the median is displayed as a line within each box. The mean values across males and females are also noted. The whiskers depict the largest and smallest data point within 1.5 * IQR. N = 7 individuals (3 male, 4 female).

### Yawns delay transitions from wake to sleep in cichlids

The occurrence of yawns in male *B. microlepis* showed clear circadian differences, with more yawns at night and dawn, when these diurnally active fish are known to rest^31^. Yawns have previously been linked to state transitions in mammals, birds, and fish^1–9^. Yawns may occur more often after waking up, or just before falling asleep, and in both cases are associated with increasing arousal. This led us to explore whether yawns were associated with state transitions in *B. microlepis*. To test this we compared the timing of behavioural transitions to the occurrence of each of the yawns in both female and male fish. To determine whether yawns occur more frequently around behavioural transitions we adapted a method used in primate species^4^. Specifically, for each yawn, we identified a matched time point one day after to be used as a time-matched control (MC) absent of yawning (“non-yawns”). We calculated the amount of time between each yawn and the next closest behavioural transition to rest. We hypothesized that if yawns function to stimulate arousal, transitions to rest should be delayed following a yawn. Our results suggest that yawns can stimulate arousal in male *B. microlepis* only. We see that state changes from active to rest (going to sleep) are less likely to occur within 30 seconds of a yawn at night compared to time matched controls (**Fig. 3a**; *P* = 0.043). This was only observed in males at night, not during the day, dawn, or dusk, or at any time in females (**Fig. S1**). In line with this, we observe that behavioural transitions are significantly delayed relative to yawns at night compared to females, whose yawns are on average ∼80 seconds closer in time to transitions (**Fig. 3b**; *P* = 5.49e-28). Because yawn frequency and timing were associated with inducing arousal in males, we next sought to determine if there was any difference in the timing or total amount of sleep between males and females. To test this we disaggregated previously published^31^ data on sleep in these fish by sex. This analysis revealed that male *B. microlepis* slept significantly less than female *B. microlepis,* which was driven by significantly less sleep specifically during the night and dawn in males (**Fig. 3c**; *P_Nigh_*_t_ = 0.016, *P_Dawn_* = 0.02). We also observe strong (*R* > 0.85; not significant) positive correlations between the number of yawns per individual and total sleep in males across time, whereas we see weaker negative correlations in females, except at night (**Fig. S2)**. At night, females show a strong negative correlation between number of yawns and total sleep (*R* = -0.97, *P* = 0.029). In males, we see a significant positive association between number of yawns and body size (*R* = 1.00, *P* = 0.0045) (**Fig. 3d**). During the dawn, we also see that the duration of male yawns is significantly positively correlated with the time of the next transition from active to rest (*R* = 0.40, *P* = 0.004) and significantly negatively correlated with y-position (*R* = -0.31, *P* = 0.027), but this was not seen for female yawns (**Fig. 3e**; *P_Male_* = 0.004, *P_Female_* = 0.98). This result is consistent with longer yawns being associated with dissipating more sleep pressure, or inducing more arousal, than shorter yawns. Overall, our results suggest that sexually dimorphic differences in yawns and behavioural transitions are associated with differences in sleep between male and female cichlid fish.

**Figure 3.**
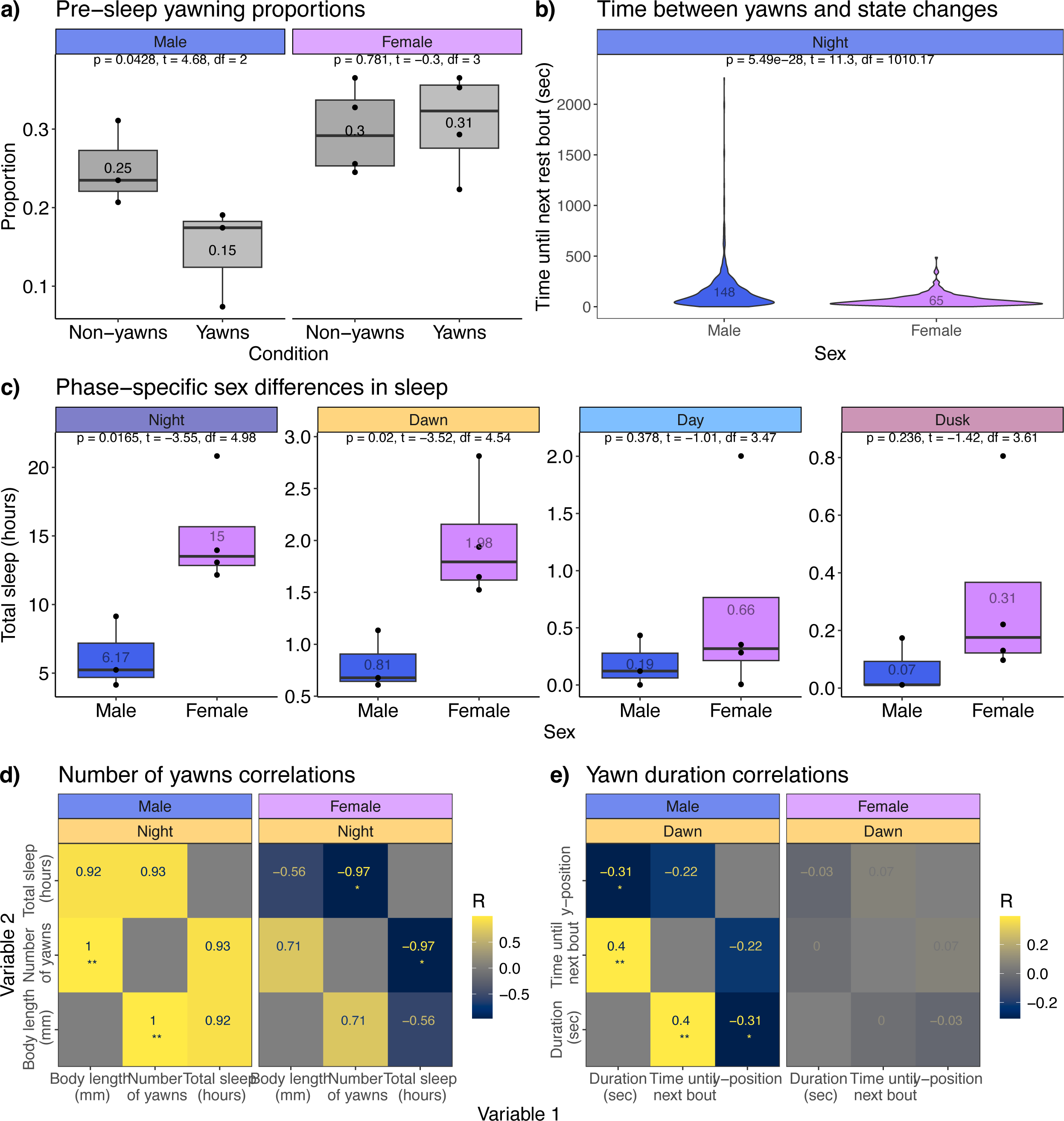
Yawns delay sleep-wake transitions in males only. **a)** Boxplots showing the proportion of night yawns occurring within 30 seconds of a behavioural transition from active to rest compared to time matched control non-yawn times. A paired t-test was used to determine the differences between conditions. **b)** Violin plot showing the time between yawns (1494 yawns; 825 male, 669 female) and behavioural transitions from active to rest for males and females at night. An unpaired t-test was used to determine differences between sexes. **c)** Boxplots showing sex differences in the total amount of sleep between males and females, split by phase. Boxes represent the interquartile range (IQR) from the first to third quartile, where the median is displayed as a line within each box. The mean values are also noted. The whiskers depict the largest and smallest data point within 1.5 * IQR. **d-e**) Correlograms showing the relationships between the total number of yawns per individual, total sleep and body size at night (**d**), and between yawn duration, time until next state change to rest, and y-position at dawn (**e**). The correlation coefficient (R) is noted in each tile. *P < 0.05, **P < 0.01. N = 7 individuals (3 male, 4 female).

### Yawns show rapid social contagion in cichlid fish

In line with the result that yawns at night and dawn are associated with state transitions, we observed that individual fish were significantly lower in the tank when yawning at these times compared to day and dusk, as seen by their mean y position during yawns in these periods (**Fig. 4a**; *P* = 6.5e-14). Furthermore, the overall yawn position density overlapped with sleep position density for each fish, which primarily occurs when the fish are at or near the bottom of the tank (**Fig. S3**). However, where yawns didn’t occur coincident with sleep, they tended to occur when the fish was nearest to the mesh divider of a neighbouring tank (**Fig. 4b**). We therefore hypothesized that these yawns could represent ‘social’ or contagious events.

**Figure 4.**
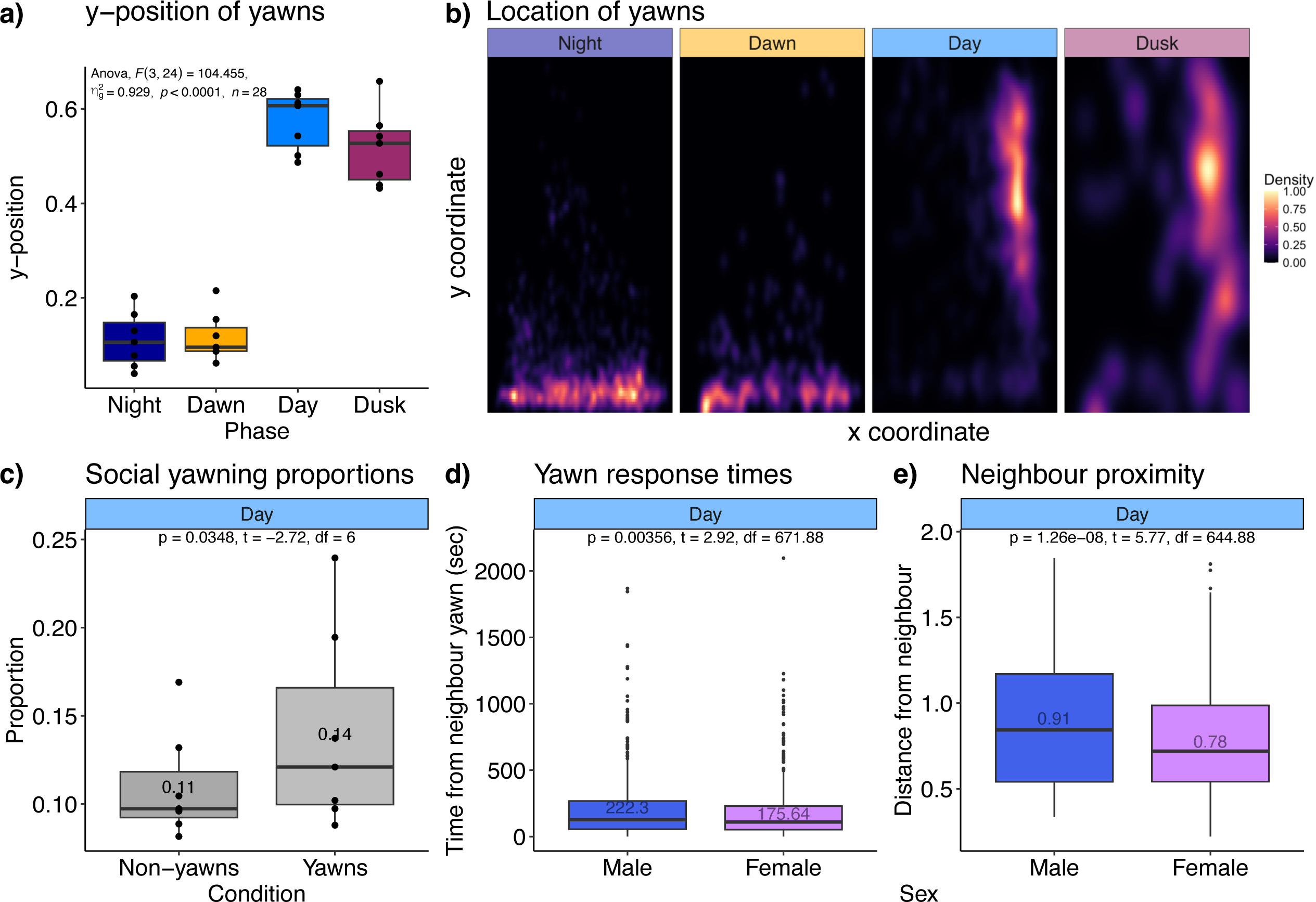
Cichlid yawns show rapid social contagion. **a)** Boxplot showing the mean y-position of yawns at different times for all individuals (males and females). An analysis of variance (ANOVA) was used to compare y-position across time between the four phases. **b)** Normalized location density for all individuals, separated by time of day. Location data have been adjusted so that in all cases the neighbour is to the right. **c)** Boxplot showing the proportion of all daytime yawns occurring within 30 seconds of a neighbour yawn compared to matched control times. A paired t-test was used to determine differences between conditions. N = 7 individuals (3 male, 4 female). **d)** Boxplots showing the time between yawns and neighbouring fish yawns during the day between males and females. **e)** Boxplot showing the mean distance between fish at the time of a daytime yawn (1019 yawns) between males and females. An unpaired t-test was used to determine differences between males and females. Boxes represent the interquartile range (IQR) from the first to third quartile, where the median is displayed as a line within each box. The mean values are also noted. The whiskers depict the largest and smallest data point within 1.5 * IQR. Outliers are shown in dots (**d - e**).

Using the same time-matched control method that we used to test for associations with state-transitions, we found that for daytime yawns (including both male and female yawns), there was an increase in the proportion of yawns occurring within 30 seconds of a neighbour yawning, compared to the time matched control (**Fig. 4c**; *P* = 0.035). This effect is only seen for yawns during the day and not the night, dawn, or dusk periods (**Fig. S4a**). Though we found no differences between sexes in the likelihood of either triggering or responding to a contagious yawn (**Fig. S4b**), compared to males, females responded with a significantly reduced delay (∼45 seconds faster on average during the day, **Fig. 4d**; *P* = 0.0036), and yawned while significantly closer in proximity to their neighbour during the day (∼1.17x closer, **Fig. 4e**; *P* = 1.26e-8). This was not observed for matched control non-yawn timepoints, where males and females did not display significant differences in their average distance from their neighbour in the absence of yawns (**Fig. S4c**). These results suggest that yawns can trigger rapid contagion in cichlid fish. Altogether, our results suggest that the social function of yawning is stronger in females, while the sleep-wake function of yawning is stronger in males.

## DISCUSSION

Yawning is a conserved and ancient behaviour, yet its function remains enigmatic across species. Yawning has been associated with the sleep-wake cycle and social cohesion through its contagious effects, and sexually dimorphic differences in yawn frequency and functions have been documented in several species^1,6–9,17,22–24,36,37^. These observations have led to three dominant hypotheses for the function of yawning, which are the state change hypothesis, the social hypothesis and the sexual dimorphism hypothesis^4,38^. In this study, we re-analyze video recordings of freely behaving and interacting *B. microlepis* as a model to assess if these three hypotheses are true for fish. Our data support the hypothesis that yawns stimulate arousal, aligning with the state change hypothesis. We also demonstrate the existence of yawn contagion in freely behaving fish. We find that yawns at night serve functions related to the sleep-wake cycle, while daytime yawns are socially-driven, supporting the social hypothesis. Importantly, we find sexual dimorphisms for both functions of yawning, providing a novel framework for the sexual dimorphism hypothesis.

Yawns in *B. microlepis* suppress transitions from active to rest, in line with yawns’ previously reported function to stimulate arousal. However, this was only seen in male individuals, where their yawn durations also impacted the timing of transitions. To our knowledge, this is the first report of sex-specific differences in the effects of yawning on the sleep-wake cycle. We suggest that this is related to the sex differences in sleep itself, where males are observed to sleep less than females. Across cichlid species, females are often the sole providers of parental care^39^. Biparental care is also commonly found, with males and females taking on different roles, often with females tending to the eggs/fry while males guard and defend the territory^39^. Breeding strategies like these would be consistent with sex differences in sleep, such that one partner has different responsibilities allowing for more/less sleep. However, it is known that the species used in the present study, *B. microlepis,* form monogamous pairs and display biparental guarding of territories and eggs^39–41^. By the fry stage of development, the roles of males and females in caring for their young are indistinguishable^41^. Differences in yawning behaviour has been reported in the cichlid *Tilapia melanotheron*, on the basis of both size and sex^42^, and body size is correlated with total sleep across species^43^. Adult individuals of *B. microlepis* are sexually dimorphic in size, and yawn frequency was associated with body size in the present study^41^. However, the individuals in our study were adolescent, therefore, expanding our analyses of yawns in different contexts and across additional cichlid species that display variations in sexual dimorphism for parental care, body size, and sleep traits could clarify these issues.

Outside of sleep-wake related yawns, we observe a high frequency of yawns during the day when the fish are near the neighbouring arena. This aligns with the social yawn hypothesis, which is built on the phenomenon of yawn contagion. Yawning is a powerful contagious behaviour, commonly explained by the need for social communication between animals^24^. Yawning has been linked to social structure in many sexually dimorphic animals, and many other social behaviours like empathy^36,44,45^, aggression^37^, and anxiety. Yawns have been observed in social contexts in cichlids^42,46^, but the contagious nature of yawning has never been directly examined in this clade. Contagion has recently been observed in zebrafish, where zebrafish were more likely to respond to a video recording of a yawning fish compared to a control video of a fish that wasn’t yawning^27^. Here, we show rapid yawn contagion, on the scale of seconds, between interacting conspecifics for the first time in fish. This expands our knowledge of yawn contagion beyond mammals, providing more insight into the evolution of this behaviour.

Female *B. microlepis* responded to neighbour yawns faster and while in closer proximity than males. However, our study design precluded thorough analysis of the contagiousness of yawns between heterologous (male-female) and homologous (same-sex) pairs. There is conflicting evidence regarding sex differences in contagious yawning in the literature. Some species displaying sexual dimorphism show differences in the amount of contagious responses. For instance, female geladas respond more to females than males^21^, and female wolves show a shorter reaction time to yawns compared to males^22^. This result is often attributed to empathy, such that females are more responsive to social cues than males^21,47^. This has been supported by the observation that individuals with closer relationships (kin and friends) show higher rates of yawn contagion and latency in humans and other animals^44,48^. Yawning is commonly described as a threat signal from males in sexually dimorphic primates^4^, and may also serve to increase vigilance within groups to prepare for potential threats^37^. In cichlid fish, many species have been observed to attack and prey on *B. microlepis* young^49^. It is possible that individual *B. microlepis* use yawns as threat displays towards conspecifics or other fish while defending their territory. However, we find that male yawns were predominantly related to the sleep-wake cycle, rather than reciprocal display yawns. Future work could assess the aggressive nature of yawning by analysing male-male or interspecies interactions.

Cichlid fish are a powerful model to probe behaviour and identify differences in closely related species. Previous work has shown cichlids occupy different temporal niches, and display remarkable diversity in sleep behaviour, including the timing and duration of sleep^31^. Our study is the first to examine yawns in the context of both the sleep-wake cycle and social behaviour in the same fish. Our results suggest that yawns can have dual functions, aligning with both the state-change hypothesis and the social hypothesis, in a sexually dimorphic manner. The complexity of yawn patterns identified here suggest that cichlids are an ideal model to study the function and evolution of yawns. In the future, comparisons could be made between social and non-social species, nocturnal (night-active) or diurnal (day-active) species, and those that differ in their sleep behaviours. Furthermore, connections could be made between yawning patterns and the diverse ecologies and morphologies present across cichlid species.

## METHODS

### Tracking of behaviour and yawning in cichlids

A subset of the data for *Boulengerochromis microlepis* (*N* = 7) processed by Nichols et al., 2025^31^ were re-analyzed for the present study. Briefly, these data were collected at the facilities of the University of Basel at the Zoological Institute. During recording, fish were housed in tanks with 24–25 °C water temperature, under 12 h of light, 12 h of darkness, with 30-min ramping light conditions at dawn (7am) and dusk (7pm). Each tank measured 45 cm height × 110 cm length × 25 cm depth, with opaque glass on three sides not facing the camera. Each tank had a thin layer of sand and was physically divided into 3 arenas (33 cm wide) which housed individual fish (**Fig. S5**). The dividers were made of PMMA opal white with a mesh insert allowing for water flow and visual, olfactory, and aural communication between fish during each tracking session. The feeding schedule was consistent with that of the home tanks, which allowed the effects of feeding to be averaged out over the six recording days. Dawn and dusk were classified as the hours before and after the light transition occurred (06:00 – 08:00 and 18:00 – 20:00). The tanks were backlit with a panel of infrared LED lights which were diffused by the opaque glass and a diffuser to generate even lighting and allow recording during the night. Fish were tested as juveniles due to the long life cycles and lack of availability of adults of these larger species. Both male (*N* = 3) and female fish (*N* = 4) were tested.

### Quantifying yawns

The first two of the six recorded days (48 hours) were analyzed for yawn quantification. A yawn was defined as when the fish opened their mouth, stretched their jaw fully (the widest point of the yawn), then closed their mouth (**Fig. 1a**). The timestamp of each yawn was recorded as the midpoint of the yawn. For a subset of yawns occurring at each time of day (1 hour of footage per phase, 327 yawns total), the duration was calculated from the start to the end of the yawn. The timing of yawns were compared to the times of the nearest rest bout to determine the time between state changes. Rest was defined as less than 5% movement less than 15 mm/sec in a rolling 60 second window^31^. Any yawn occurring while the fish was defined as resting was excluded.

### Statistics and reproducibility

All analyses were conducted in R (v4.4.1). Paired t-tests were used for comparisons between test and control groups, and unpaired t-tests were used for sex comparisons. A Shapiro-Wilk test was used to determine that both the sleep-wake yawning proportions (*P* = 0.73) and social yawning proportions (*P* = 0.16) were normally distributed. An analysis of variance (ANOVA) was used for phase comparisons. To normalize the number of yawns at each phase, the total number of yawns was divided by the total number of hours spent in each phase (4 h for dawn and dusk, 10 h for night and day). Plots were made in R using packages ggplot2 (v4.0.0)^50^, ggpmisc (v0.7.0)^51^, ggpubr (v0.6.3)^52^, ggh4x (v0.3.1)^53^, ggnewscale (v0.5.2)^54^, patchwork (v1.3.2)^55^, and viridis (v0.6.5)^56^. The distance (d) between two fish was calculated using the formula 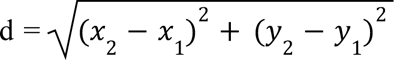, where x_1_ and y_1_ represent the coordinates of one fish and x_2_ and y_2_ represent the coordinates of the neighbouring fish. Point patterns for the locations of yawns and sleep were generated using spatstat (v3.6.0)^57^. Additional libraries used for analysis include lubridate (v1.9.5)^58^, tidyr (v1.3.2)^59^, dplyr (v1.2.1)^60^, broom (v1.0.12)^61^, gsheet (v0.4.6)^62^, tictoc (v1.2.1)^63^, data.table (v1.18.2.1)^64^, stringr (v1.6.0)^65^.

To determine whether yawning was more likely to occur before or after behavioural transitions, we adapted a method previously developed for post-conflict behaviour in primates and later used in yawn behavioural studies^4^. Each observed yawn was compared to a time matched control timepoint one day after the observation. The matched control time was checked to ensure that an actual yawn was not observed at the time. Only two yawns were observed at the same time of the time matched control. The closest behavioural transition from active to rest was calculated for both the observed yawn and MC non-yawn. Then, the proportion of yawns that were within 30 seconds of a behavioural transition were compared between the yawns and matched controls. Given the highly fragmented nature of fish sleep^35^ (Mean rest bout length = 79 seconds), we used a shorter interval of 30 seconds to better capture the relationship between yawns and rest.

The same method was used to determine whether neighbouring fish were more likely to yawn sequentially (following another observed fish yawn). Here, the time matched control was either one day after or one day before the observed yawn, depending on the day. If the yawn occurred on the first day, the time MC observation was on the next day, while yawns occurring on the second day were matched to the previous day. The proportion of yawns that were within 30 seconds of an observed neighbour yawn were compared between the yawns and matched controls.

## Data and code availability

Source code used in this study can be assessed on GitHub: https://github.com/ayashaa/yawning_analysis.

## ACKNOWLEDGEMENTS

We thank members of the Shafer laboratory for discussion and advice and A. Nichols and M.B. Sokolowski for valuable comments on this manuscript. We thank the STEM Research Fellowship Research Exploration Opportunity (REO) program for providing the framework to allow for the participation of V.K., R.K, J.S., P.M., C.S., and S.R. in this project. This work was supported by grants from the National Sciences and Engineering Research Council of Canada (NSERC) (RGPIN-2024-05509) and the Swiss National Science Foundation (SNSF) to M.E.R.S. (196313), and a Canada Graduate Research Scholarship from NSERC to A.A.W.

## AUTHOR CONTRIBUTIONS

A.A.W and M.E.R.S. conceived and designed the study. V.K., R.K, J.S., P.M., C.S., and S.R. collected behavioural data. A.A.W collected and analysed behavioural data. A.A.W. and M.E.R.S. interpreted the results and wrote the manuscript. All authors read and approved the manuscript.

## SUPPLEMENTAL FIGURES

**Figure S1.**
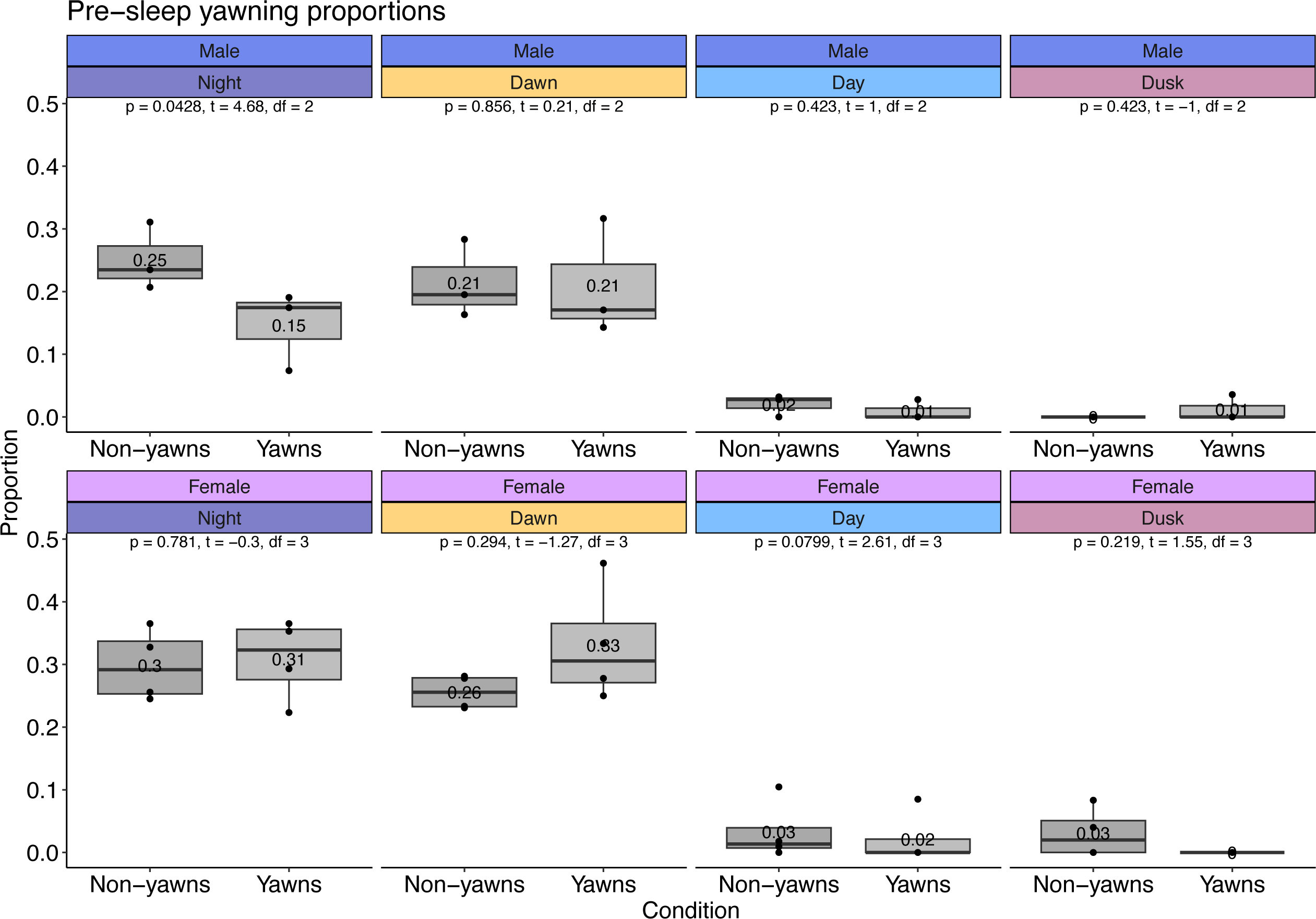
Yawns delay sleep-wake transitions at night in males only. Boxplots showing the proportion of yawns occurring within 30 seconds of a state change from active to rest compared to matched control non-yawn times, separated by both phase and sex. A paired t-test was used to compare conditions. Boxes represent the interquartile range (IQR) from the first to third quartile, where the median is displayed as a line within each box. The mean values are also noted. The whiskers depict the largest and smallest data point within 1.5 * IQR. N = 7 (3 male, 4 female).

**Figure S2.**
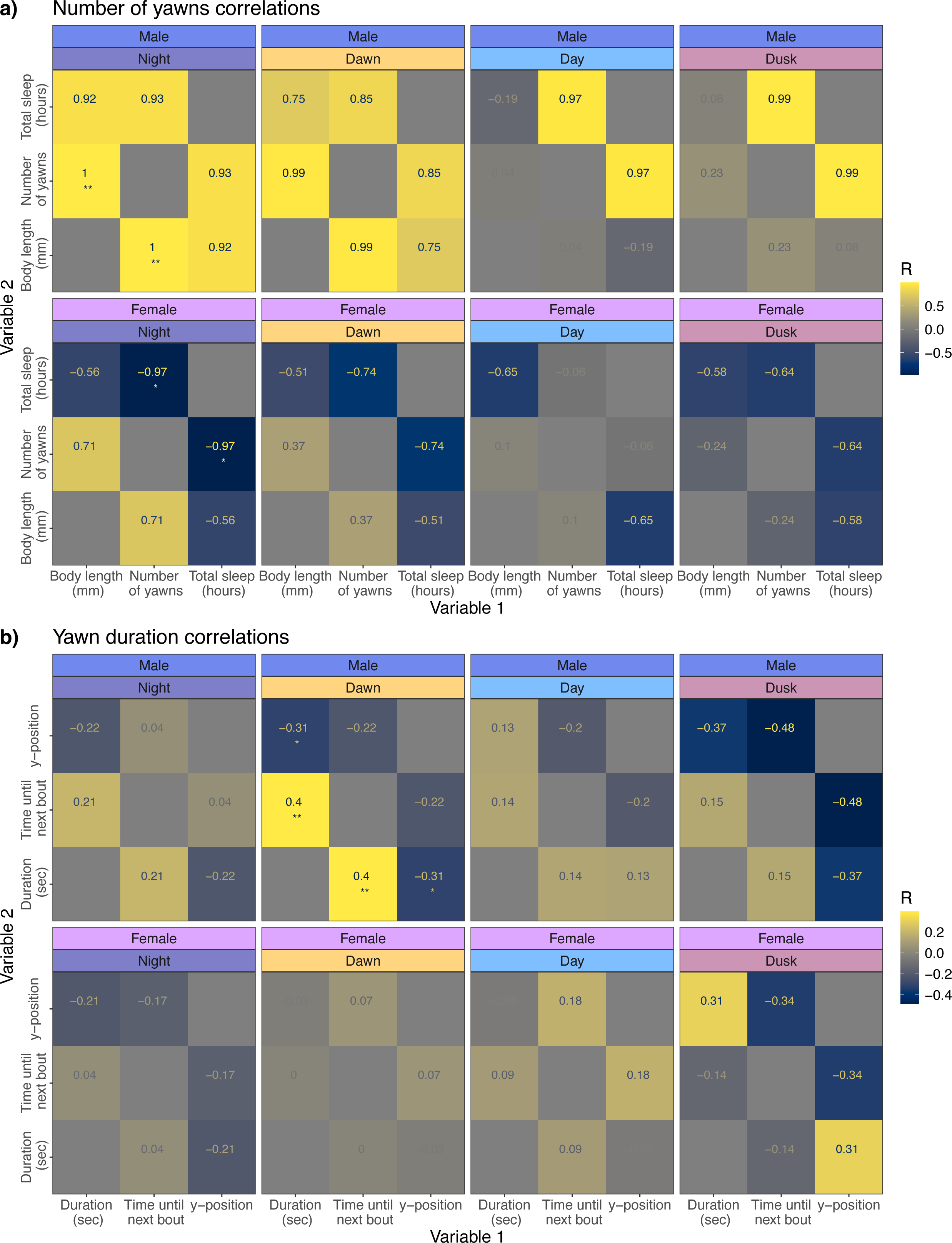
Yawning behaviour is correlated with body size and state changes in males only. **a-b**) Correlograms showing the relationships between the total number of yawns per individual, total sleep and body size (**a**), and between yawn duration, time until next state change to rest, and y-position (**b**). Split by both sex and time of day. The correlation coefficient (R) is noted in each tile. *P < 0.05, **P < 0.01. N = 7 (3 male, 4 female).

**Figure S3.**
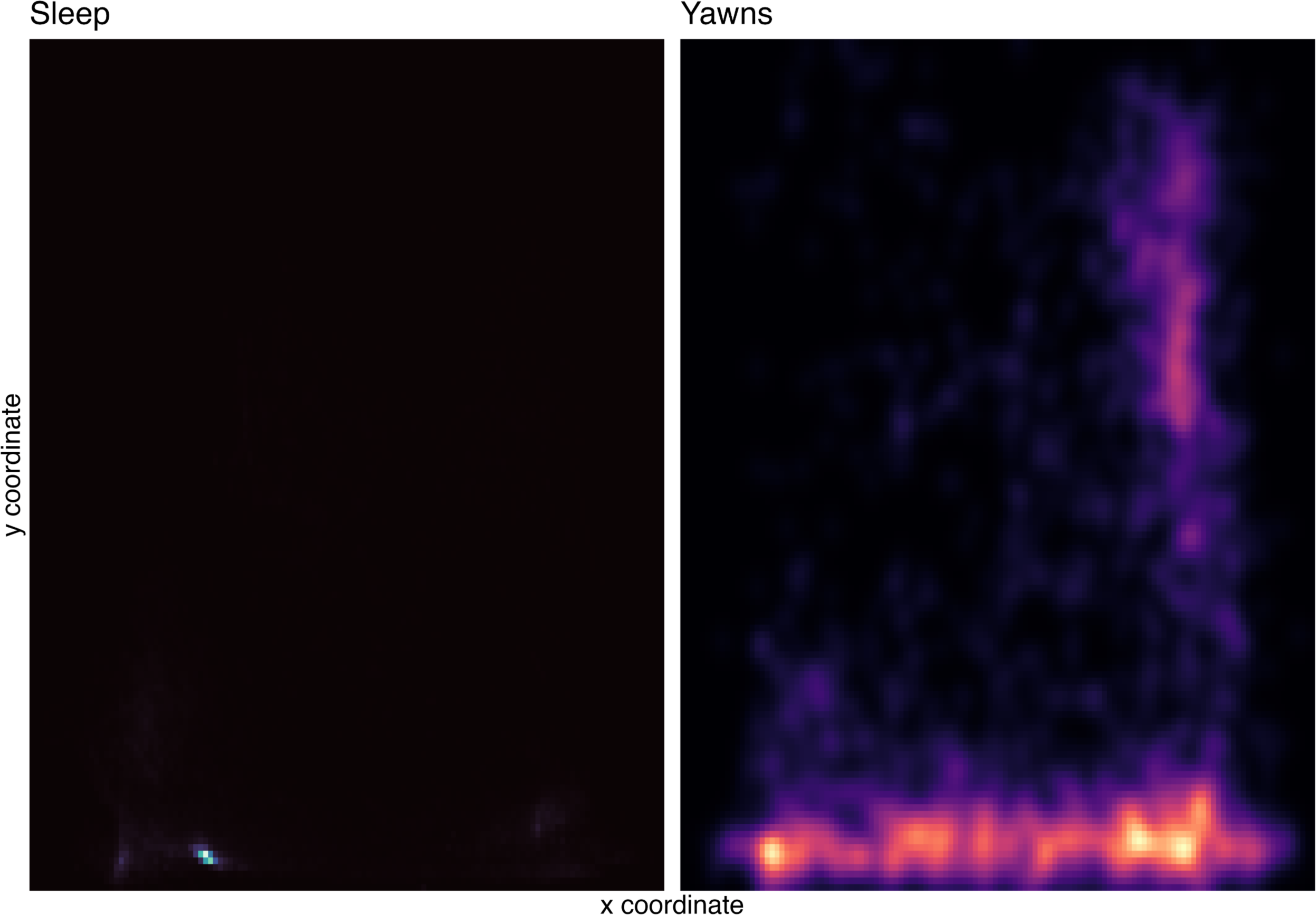
Sleep and yawn locations overlap. Normalized location densities for sleep and yawns, combined across all individuals tested.

**Figure S4.**
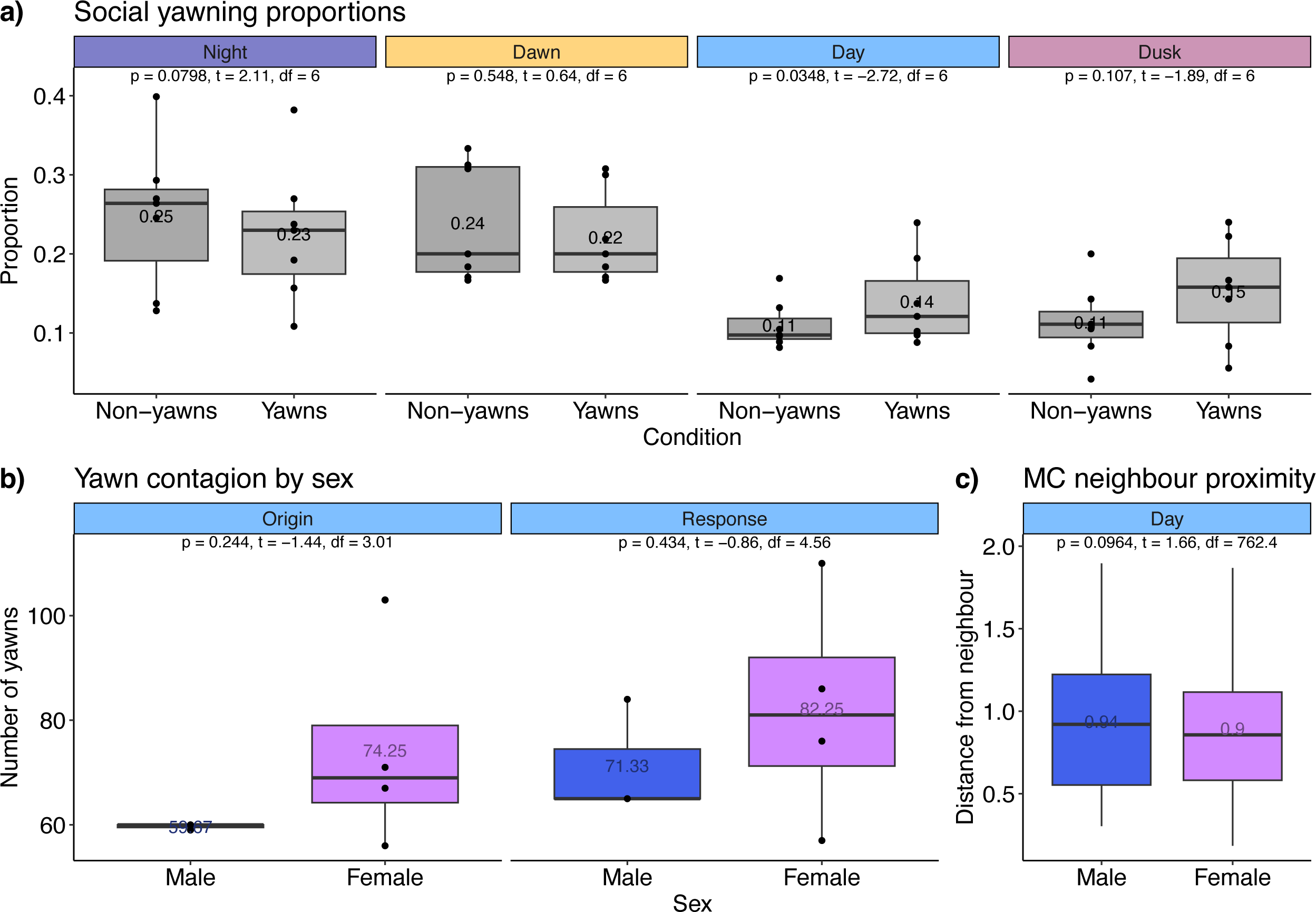
Yawns in cichlids show contagion during the day only. **a)** Boxplots showing the proportion of yawns occurring within 30 seconds of a neighbour yawn compared to matched control (MC) non-yawn times, separated by phase. A paired t-test was used to compare conditions. N = 7. **b)** Boxplots showing the number of yawns that were the origin or response of contagious yawn events during the day. **c)** Boxplot showing the mean distance between fish at the MC non-yawn times during the day between males and females. An un-paired t-test was used to compare males and females. N = 1029 daytime yawns. Boxes represent the interquartile range (IQR) from the first to third quartile, where the median is displayed as a line within each box. The mean values are also noted. The whiskers depict the largest and smallest data point within 1.5 * IQR.

**Figure S5.**
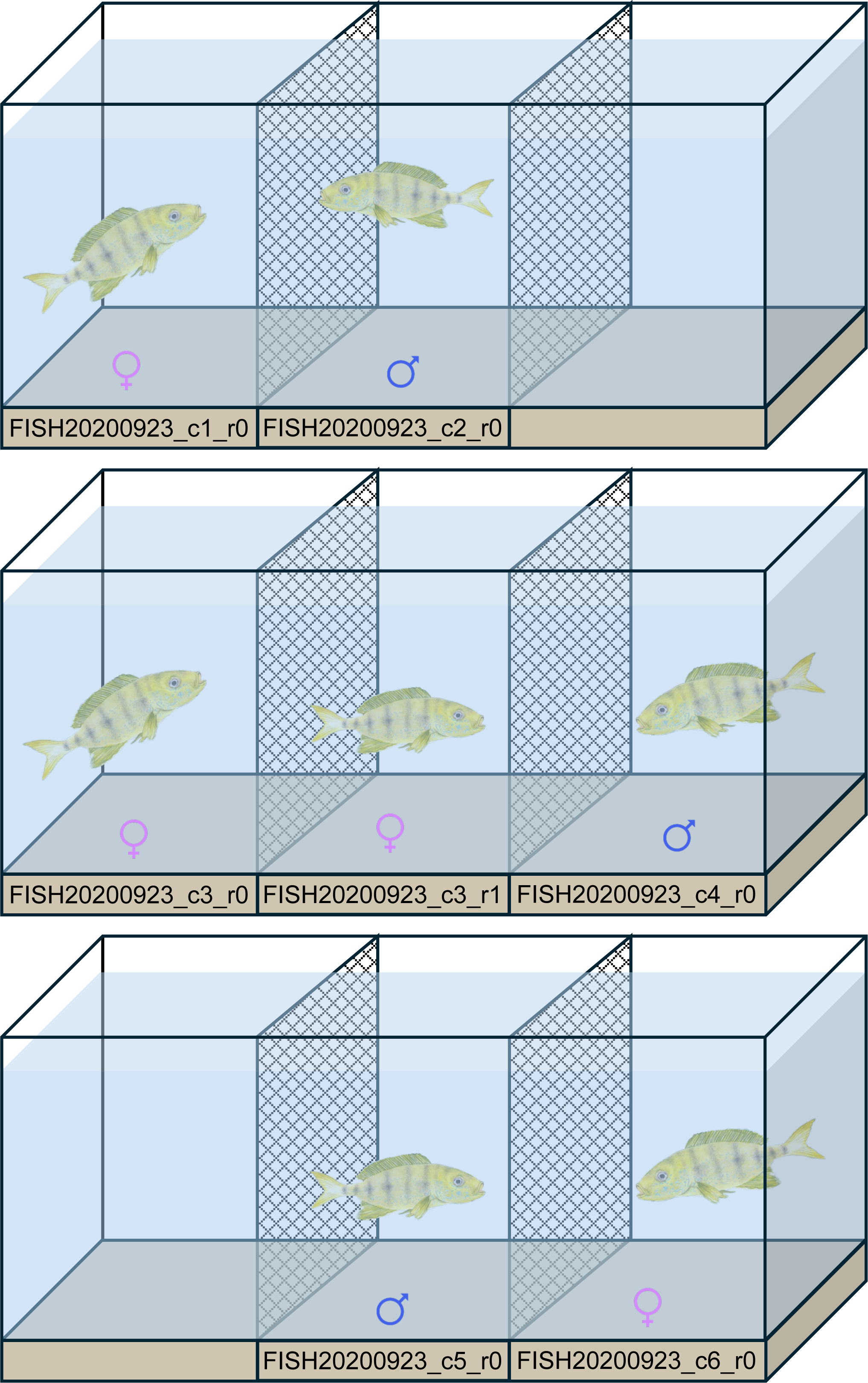
Diagram of tank setup. Fish were housed in tanks with 2-3 individuals, separated by mesh dividers.

